# Matrix Inversion and Subset Selection (MISS): A novel pipeline for mapping of diverse cell types across the murine brain

**DOI:** 10.1101/833566

**Authors:** Christopher Mezias, Justin Torok, Pedro D. Maia, Eric Markley, Ashish Raj

## Abstract

The advent of increasingly sophisticated imaging platforms has allowed for the visualization of the murine nervous system at single-cell resolution. However, current experimental approaches have not yet produced whole-brain maps of a comprehensive set of neuronal and nonneuronal types that approaches the cellular diversity of the mammalian cortex. Here we aim to fill in this gap in knowledge with an open-source computational pipeline, Matrix Inversion with Subset Selection (MISS), that can infer quantitatively validated distributions of diverse collections of neural cell types at 200μm resolution using a combination of single-cell RNAseq and *in situ* hybridization datasets. We rigorously demonstrate the accuracy of MISS against literature expectations. Importantly, we show that gene subset selection, a procedure by which we filter out low-information genes prior to performing deconvolution, is a critical pre-processing step that distinguishes MISS from its predecessors and facilitates the production of cell type maps with significantly higher accuracy. We also show that MISS is generalizable by generating high-quality cell type maps from a second, independently curated single-cell RNAseq dataset. Together, our results illustrate the viability of computational approaches for determining the spatial distributions of a wide variety of cell types from genetic data alone.

## INTRODUCTION

Characterizing whole-brain distributions of neural cell types is a topic of keen interest in modern neuroanatomy, with many applications to both basic and clinical neuroscience research (Arneson et al., 2018; Fu et al., 2018, 2019; Muratore et al., 2017; Rama Rao et al., 2018; Skene et al., 2018). Advances in molecular methods for quantifying gene expression and data analytic cell clustering techniques based on morphologic or genetic profiles are enabling the mapping of meso- and microscale neuronal and non-neuronal cell type architecture at a whole-brain level (Codeluppi et al., 2018; Erö et al., 2018; Kim et al., 2017; Moffitt et al., 2018; Murakami et al., 2018; Tasic et al., 2018; Wang et al., 2018; Zeisel et al., 2018). Mammalian whole-brain cell type mapping has historically focused on neuromodulatory systems, largely because the identification of catecholamine-producing subpopulations using molecular markers is rather straightforward (Beier et al., 2015; Björklund and Dunnett, 2007; Pazos et al., 1985; Vincent and Kimura, 1992). More recently, serial two-photon tomography (STPT) imaging of cells expressing individual cell type markers genetically tagged to green fluorescent protein (GFP) successfully mapped three important subpopulations of inhibitory GABAergic interneurons (Kim et al., 2017) and cholinergic neurons (Li et al., 2017) across the murine brain. Although the above animal laboratory techniques cover the entire brain, they are expensive and time-consuming to apply to mice, impractical to apply to larger-brained model organisms, and impossible to apply to human subjects.

Recent computational work demonstrates the feasibility of using existing datasets of cell type gene expression or cell markers for mapping cells in space across the vertebrate brain. Recent work produced whole-brain maps of broad classes of neuronal and glial subpopulations at single-cell resolution using purely computational methods (Erö et al., 2018), with the limitation that the mapped cells were classed into large meta-groups, such as GABAergic versus glutamatergic neurons, rather than more specific cell types. Pioneering work mapping single-cell RNA sequencing (scRNAseq) data from aquatic flatworms and zebrafish onto *in situ* hybridization (ISH) expression provided a plausible route to mapping highly specified cell types in mammalian nervous systems (Achim et al., 2015; Satija et al., 2015). Others have inferred the spatial distribution of cell types by deconvolving type-specific microarray expression profiles from spatially realized ISH expression data (Grange et al., 2014). In theory, using a matrix-inversion-based approach provides a better estimate of cell density than correlation-based mapping, because cell types with highly similar gene expression profiles will necessarily have highly correlated spatial profiles. However, the original work pioneering matrix inversion for cell type mapping provided no external quantitative validation of their maps. We hypothesized that, with several modifications, the approach of deconvolving ISH data into cell type densities using recently available scRNAseq data could create voxel-level cell type maps that could be externally validated both qualitatively and quantitatively.

We present an information-theoretic computational pipeline that can supply per-voxel estimates of cell densities of distinct and highly specified cell types across the whole murine brain, which is the best characterized mammalian nervous system at a molecular level. After deriving matrices of type-specific expression profiles from scRNAseq data (AIBS, 2018; Tasic et al., 2018; Zeisel et al., 2018) and the matrix of spatial gene expression information from the Allen Institute for Brain Science (AIBS) ISH atlas (Lein et al., 2007), we then solve a linear system of equations for cell type density per voxel using a nonnegative least-squares algorithm, like prior approaches (Grange et al., 2014). Two key methodological innovations make the present maps possible. First, we hypothesize mapping accuracy will improve dramatically if we filter out all “low-information” genes prior to matrix inversion. Although using only a subset of the genes runs counter to similar prior approaches (Andersson et al., 2020; Grange et al., 2014), we suspected the inclusion of genes that are lowly expressed or poorly differential across cell types would deteriorate the quality of the resulting maps. Therefore, we developed a novel information-theoretic algorithm, **Minimal Redundancy – Maximum Relevance – Minimum Residual (MRx3)**, selecting an optimal subset of genes to capture cell densities across the mouse brain (see **Methods** for details) and used it to perform subset selection prior to matrix inversion. Second, we formulated several objective metrics to assess the biological accuracy of cell type maps, relying in part upon regional quantification of individual cell types where available (Kim et al., 2017). We further evaluated the generalizability of our approach by mapping a larger, more widely sampled, and independently collected scRNAseq dataset (Zeisel et al., 2018).

Overall, we demonstrate that both MRx3-based subset selection and matrix inversion are necessary to achieve superior cell type maps as compared with correlation-based methods and deconvolution/inversion methods without subsetting (Grange et al., 2014). All cell-type-specific maps across both scRNAseq datasets are available for download along with this methodological pipeline, which we call **Matrix Inversion with Subset Selection (MISS)**. The MISS pipeline is designed to project any arbitrary single-cell expression data onto any arbitrary spatial expression atlas and can be applied to other brains, such as those from macaques and humans. The aim of the current paper is to demonstrate the accuracy of the present maps and the importance of gene subset selection with data-driven approaches such as our MRx3 algorithm using mouse data where quality assessments are tractable.

## RESULTS

### Overview of Matrix Inversion with Subset Selection (MISS)

A schematic of the MISS pipeline is displayed in **Figure 1**. We extracted publicly available ISH (Lein et al., 2007) and scRNAseq data (AIBS, 2018; Tasic et al., 2018; Zeisel et al., 2018) and collected all overlapping genes between scRNAseq and ISH datasets (**Figure 1, Step 1**). Since nonspecific genes can introduce noise and other artifacts, we developed a novel information-theoretic algorithm for reordering the genes, referred to herein as “MRx3” (**Figure 1, Step 2**). After this reordering, we find the elbow of the residual curve, defined as the point closest to the origin, and exclude all genes past this point (**Figure 1, Step 3**). We then use the solution to the non-negative matrix inversion using only the genes included in the subset after elbow selection to yield densities per voxel (**Figure 1, Steps 4 & 5**). Notably, all 3D whole-brain illustrations of all cell types in this paper are not at individual cell resolution as our method does not produce individual cell locations. Rather, they are voxel-level point cloud illustrations (see **Methods** for details) with the density of points per voxel controlled by the density of that type of cell in that voxel. MISS-inferred densities for all cell types using scRNAseq data from the AIBS (AIBS, 2018; Tasic et al., 2018), including all neurons and glia, can be found in **S. Table 4**. This dataset, which we refer to throughout the paper as the “Tasic *et al*. dataset,” pools data collected from the visual and motor cortices as well as the lateral geniculate complex of the thalamus. See **Methods** for a full description of the MISS pipeline and the optimization procedure. We will refer to results generated the elbow selected gene set after MRx3 reordering followed by inversion as results from “MISS.” Further methodological details, such as the hierarchical clustering levels at which we map the Tasic *et al*. scRNAseq dataset and the elbow curves for mapping both scRNAseq datasets, can be found in **S. Figure 1**.

**Figure 1.**
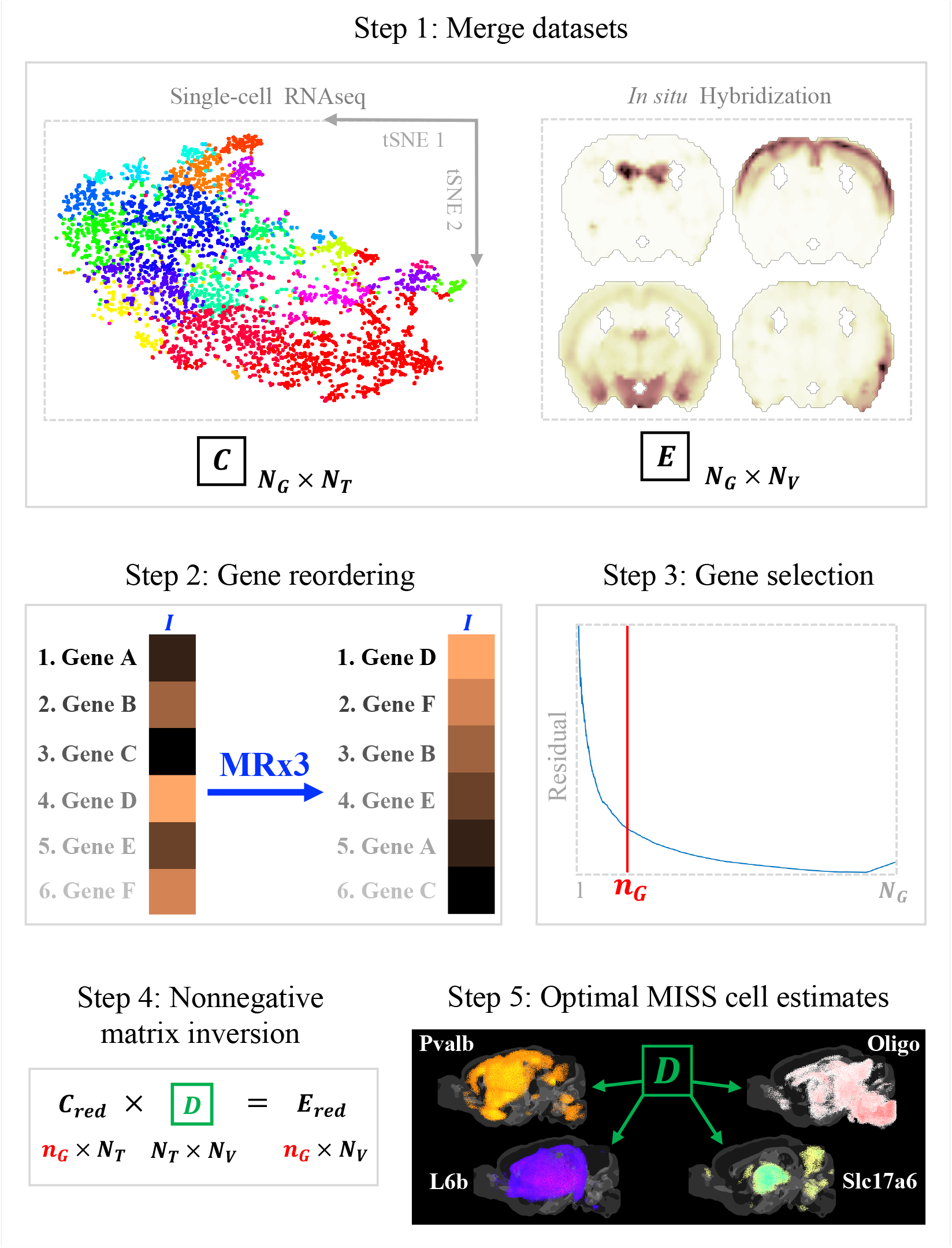
A visual outline of the MISS pipeline for mapping cell type clusters. *Step 1* consists of finding appropriate scRNAseq cell cluster expression data and combining it with spatial gene expression data, such as the AIBS gene expression atlas (Lein et al., 2007) used here. In *Step 2* the MRx3 algorithm posed here is used to reorder the genes according to information content relevant to the mapping problem at hand. *Step 3* chooses a cutoff point in the reordered gene list and only genes ranked at or above the selected index value, *n*_G_, are used for inversion. This subset selection is accomplished by plotting subset size versus the residual and then choosing an elbow, defined as the point on the curve closest to the origin. The inversion using only the chosen genes and the MISS-inferred maps produced are *Steps 4 & 5*, respectively.

### Matrix inversion with gene subset selection produces quantitatively superior maps

**Figure 2a** shows whole brain illustrations of the Tasic *et al*. MISS results, using the selected MRx3 ordered geneset, for *Pvalb*+, *Sst*+, and *Vip*+ interneurons at *n*_G_ = 606 (see **S. Figure 1b** for elbow curve). We achieve significant quantitative agreement with interneuron densities reported in prior work (Kim et al., 2017) across the neocortex, with Pearson’s R = 0.84 and Spearman’s ρ = 0.85 for *Pvalb*+ cells, R = 0.52 and ρ = 0.59 for *Sst*+ cells, and R = 0.63 and ρ = 0.66 for *Vip*+ cells (all *p* < 0.001) (**Figure 2b**). Although using inversion without performing MRx3 subset selection yielded maps of similar quality for *Pvalb*+ interneurons (R = 0.83, ρ = 0.86, *p* < 0.001), it resulted in significantly worse *Sst*+ (R = 0.38, ρ = 0.37, *p* < 0.05) and *Vip*+ interneuron maps (R = 0.24, ρ = 0.25, *p* > 0.05) (**Figure 2c**). The correlation-based mapping procedure also generally performed worse than MISS at recreating *Sst*+ and *Vip*+ interneuron distributions, but performance improved when we excluded genes that failed to satisfy the MRx3 algorithm (**Figures 2d & 2e**). Overall, our hypothesis that data-driven subset selection with matrix inversion would be important for the quality of cell type or class maps is confirmed by these interneuron results, as we can only achieve high-accuracy results for all three cell types with MISS.

**Figure 2.**
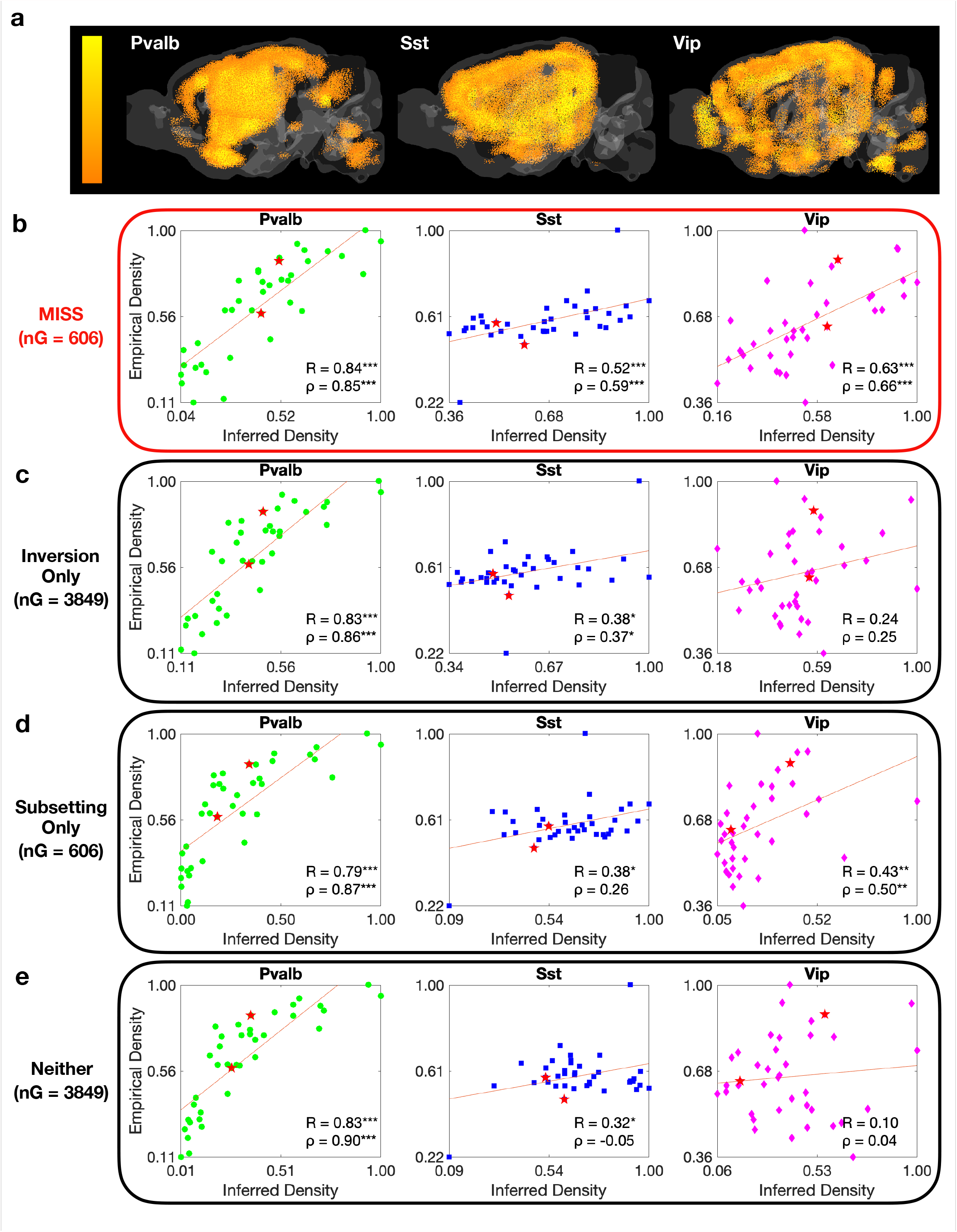
Matrix inversion after MRx3 gene subset selection produces *Pvalb*+, *Sst*+, and *Vip*+ generally outperforms inversion without subset selection and correlation-based mapping. (a) Sagittal axis views of whole brain MISS maps of *Pvalb*+, *Sst*+, and *Vip*+ interneurons in the mouse common coordinate framework (CCF) (Oh et al., 2014). Scatterplots depicting correlations between empirical measurements of *Pvalb*+, *Sst*+, and *Vip*+ interneuron densities across neocortical regions (Kim et al., 2017) and (b) MISS estimates, (c) matrix inversion without gene subsetting, (d) correlation-based mapping using the chosen MRx3 gene subset, and (e) correlation-based mapping using all the genes. Red asterisks indicate sampled regions in the scRNAseq dataset (Tasic et al., 2018). \**p* < 0.05, \*\**p* < 0.01, \*\*\**p* < 0.001.

### MISS layer-specific cell type distributions reproduce neocortical laminar architecture

We next used a metric based on Kendall’s *τ, τ*_adj_ (see **Methods** for details), to compare the ordering of laminar glutamatergic projection neurons in our maps versus their expected order given identity and sampling location (Tasic et al., 2018). Using the MRx3 gene subset at *n*_G_ = 606 (**S. Figure 1b**) with matrix inversion yields *τ*_*adj*_ = 0.75, while matrix inversion without subset selection, *τ*_*adj*_ = 0.57, and correlation-based mapping, *τ*_*adj*_ = 0.56, both perform worse (**Figure 3a**). Qualitative assessment finds that Layer-2/3 (L2/3) neurons inferred by MISS are most enriched in a band barely inside of the cortical surface. In contrast, L6 neuron enrichment forms a band that traces the interior border between the neocortex and white matter tracts, demarcated by ventricles. L4 and L5 neurons show enrichment in bands that are intermediary between L2/3 and L6 neurons, in the expected order (**Figure 3a**). Notably, maps produced with matrix inversion without subset selection appear to be worse than our optimal MISS maps because they contain more non-neocortical, and therefore off-target, estimated cell density for these types (**Figure 3a**). However, while correlation-based mapping using the subset selected gene set has very little off-target expression, akin with optimal MISS maps, it does not produce clearly defined bands for each expected cortical layer, likely explaining its lower *τ*_adj_ compared with MISS maps (**Figure 3a)**.

**Figure 3.**
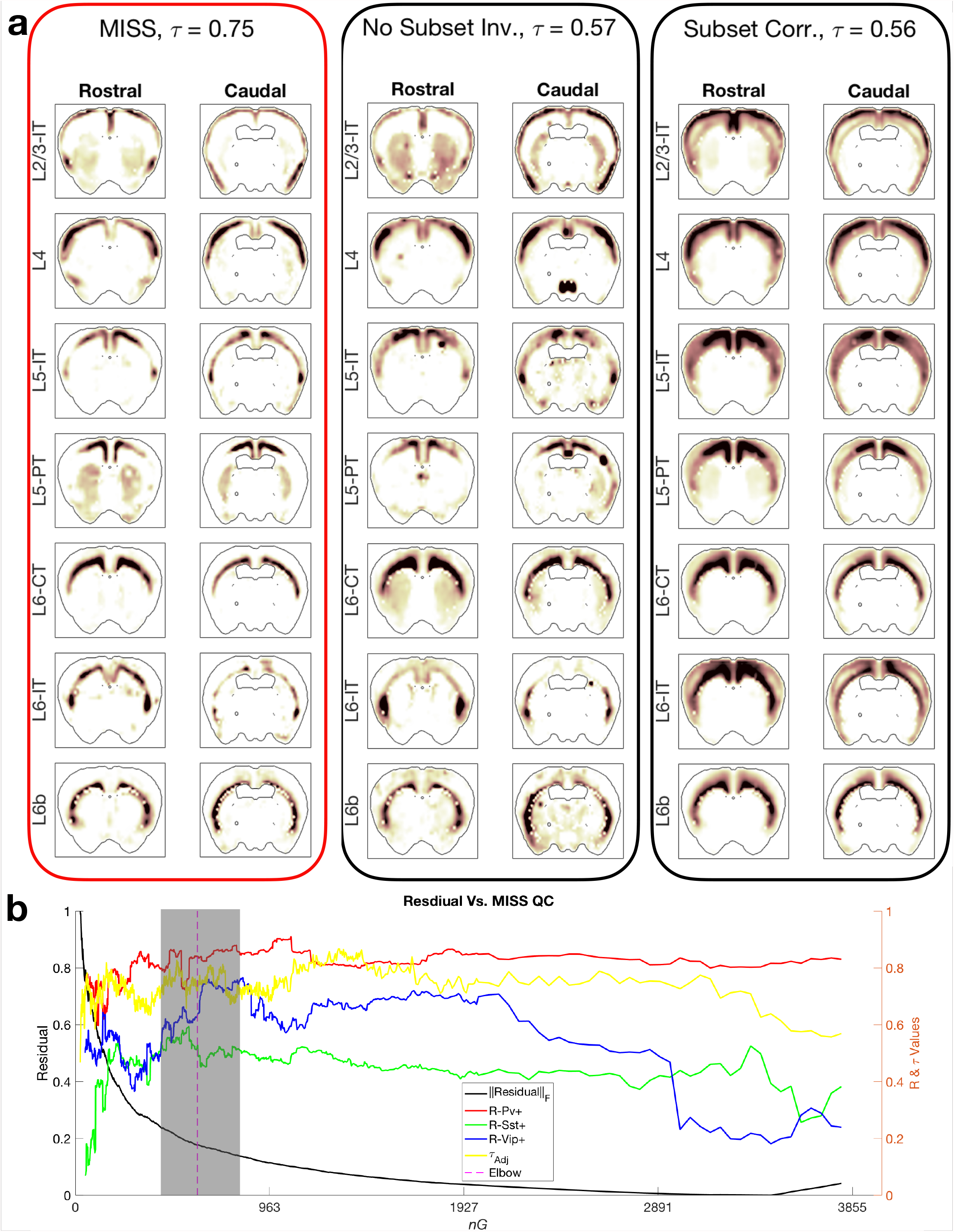
MISS with MRx3 produces laminar glutamatergic projection neuron maps that generally outperform those produced with inversion without gene subset selection and correlation-based maps. (a) The *τ*_adj_ value for MISS projection neuron maps (left) is better than inversion using all available genes (center) as well as correlation-based mapping using the MRx3-based subset (right). In general, the MISS maps produce clear bands for each projection neuron class in the appropriate cortical layer, which are less clear the correlation-based maps, while there is more significant off-target signal when there is no prior gene subset selection. (b) Within a range of about 100 genes on either side of the chosen elbow, both the interneuron R-values and the laminar excitatory neuron ordering metric *τ*_adj_ jointly achieve peak or close to peak performance, indicating that the elbow of the residual curve is a suitable ground-truth-independent metric on which we choose gene subset size, *n*_G_.

### MISS provides stable, accurate cell type maps without notable overfitting

We note that our methodological decision to choose an elbow *n*_G_ based on residual error, a ground-truth independent measure, does not produce a uniquely high-performing gene subset; **Figure 3b** shows that, relative to the elbow curve, any *n*_G_ subset size between ∼520 and ∼675 will outperform using the entire set of genes. This range of values also produces maps that correlate with those using the elbow subset (**S. Figure 2c**), indicating that our results are not a product of overfitting. Our maps show no bias towards scRNAseq sampled regions, i.e., the distribution of absolute error (in all voxels) for scRNAseq-sampled and non-sampled regions did not differ (**S. Figure 2b**). Similarly, a spatial map of each region’s average per-voxel residual did not denote highlight sampled regions in a whole-brain 3D illustration (**S. Figure 2a**). This result indicates that, at the level of cell type specificity mapped, types from the scRNAseq dataset were of types general enough to map across many brain regions without bias toward sampled regions.

### Glial cells have whole-brain distributions reflective of their biological roles

**Figure 4a** shows spatial reconstructions of the three glial cell types within the Tasic *et al*. scRNAseq dataset. Astrocytes, are most concentrated in regions of the cerebellum, the site of heaviest gray matter concentration in mammals (Keller et al., 2018; Ma et al., 2005) (**Figures 4a & 4d**). Given the relatively high density of neurons in the cerebellum and the integral support roles astrocytes play, it may be expected that astrocyte density is most pronounced in the cerebellum as well. Meanwhile, visualizations of oligodendrocytes, a glial cell type important for maintaining axon myelination (Bradl and Lassmann, 2010), expectedly trace major white matter tracts and have a marked enrichment in ventromedial regions of the brain (**Figures 4b & 4d**); indeed, at the sub-millimeter resolution afforded by the ISH atlas, the corpus callosum is clearly delineated (arrows). We also show widespread microglial distribution across much of the mouse brain (**Figures 4c & 4d**). Finally, we produce maps of sampled blood vessel cells pooled together they qualitatively resemble the major veins and arteries of the mouse brain (**Figure 4e**). Overall, voxel-level MISS maps of non-neuronal cells agree qualitatively with their expected localizations given their specialized roles. MISS inferred densities of glia can be found in **S. Table 4**.

**Figure 4.**
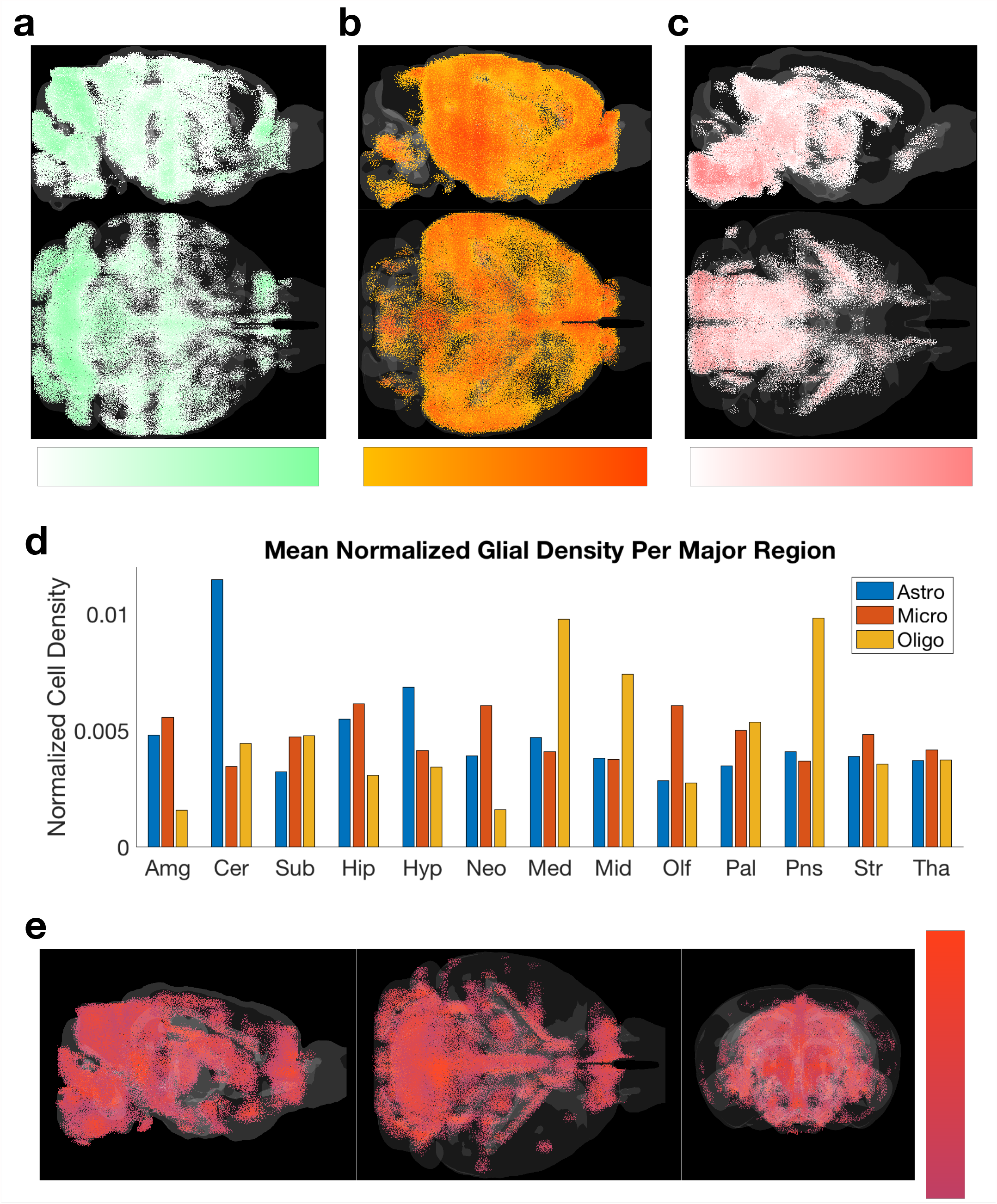
Glial and blood vessel cell maps produced using MISS generally conform with prior expectations. We show axial and sagittal views of the whole-brain maps of (a) astrocytes, (b) microglia, and (c) oligodendrocytes in CCF space. (d) Normalized densities of these glial cell classes in the projected MISS maps show astrocyte density is highest in cerebellum, oligodendrocytes are densest in the brainstem, and microglia are spread relatively evenly across the brain. (e) Three views of the maps of blood vessel cells (a combined class of endothelial and pericyte cells from the Tasic *et al*. taxonomy) in CCF space.

### MISS successfully maps cell types in an independent scRNAseq dataset

Despite our successful mapping of the cell types from the Tasic *et al*. scRNAseq dataset, it remained unclear if the MISS pipeline would generalize to other datasets, particularly those with more numerous and more finely specified cell types. Therefore, we applied MISS to the Zeisel *et al*. scRNAseq dataset, which contains 200 cell types sampled throughout the entire mouse brain (Zeisel et al., 2018). For comparison, we recreated the maps presented by the original authors using the combined set of differential genes they identified across cell types, which we then correlate at the voxel level with the output of the MISS pipeline using the MRx3-chosen gene set (elbow *n*_G_ = 1360; see **S. Figure 1c**).

We first examined the maps from four individual cell types, TEGLU12 (a telencephalic glutamatergic neuron), DEINH2 (a diencephalic GABAergic neuron), MBDOP2 (a midbrain dopaminergic neuron), and MOL3 (a type of oligodendrocyte). These cell types were selected for three reasons: 1) they represent a sampling of different general cell classes in the brain (excitatory, inhibitory, and modulatory neurons, as well as a glial cell); 2) these cells were sampled from independent regions, so 4 out of 12 sampled regions are represented here; 3) these cell types were mapped directly by Zeisel *et al*. Our MISS-derived TEGLU12 neuronal map correlates strongly with its correlation-based counterpart (R = 0.67), with expression limited to the ventral part of the inner neocortex for both (**Figure 5a**, top and center panels). Both procedures produce maps of DEINH2 neurons that indicate strong thalamic expression, and high correlation at the voxel level (R = 0.55) (**Figure 5b**). The 3D visualization of DEINH2 neurons confirms that thalamic nuclei contain by far the highest density of these cells (**Figure 5f**, 2^nd^ from the top panel). Similarly, MBDOP2 cells are concentrated in the substantia nigra for both mapping procedures, as would be expected of subtypes of midbrain dopaminergic neurons with strong spatial correlation (R = 0.59) across the whole brain (**Figure 5c**). The 3D visualization for these midbrain dopaminergic neurons also indicates predominantly nigral expression (**Figure 5f**, 3^rd^ from the top panel). Finally, MOL3 cells, a subtype of oligodendrocyte, have distributions that are both visually similar and highly correlated with an R-value of 0.63 (**Figure 5d**). MISS maps for MOL3 oligodendrocytes indicate a caudal bias consistent with the maps from Zeisel *et al*. (**Figure 5f**, bottom panel). Generally, we found the two sets of maps were highly correlated across all types within the several major classes of cell types contained in this dataset (**Figure 5e**), with overall median and mean R values of 0.56 and 0.54, respectively, at the per-voxel density level. Taken together, despite significant differences in the protocols for mapping, these results demonstrate that MISS faithfully reproduces expected cell type distributions.

**Figure 5.**
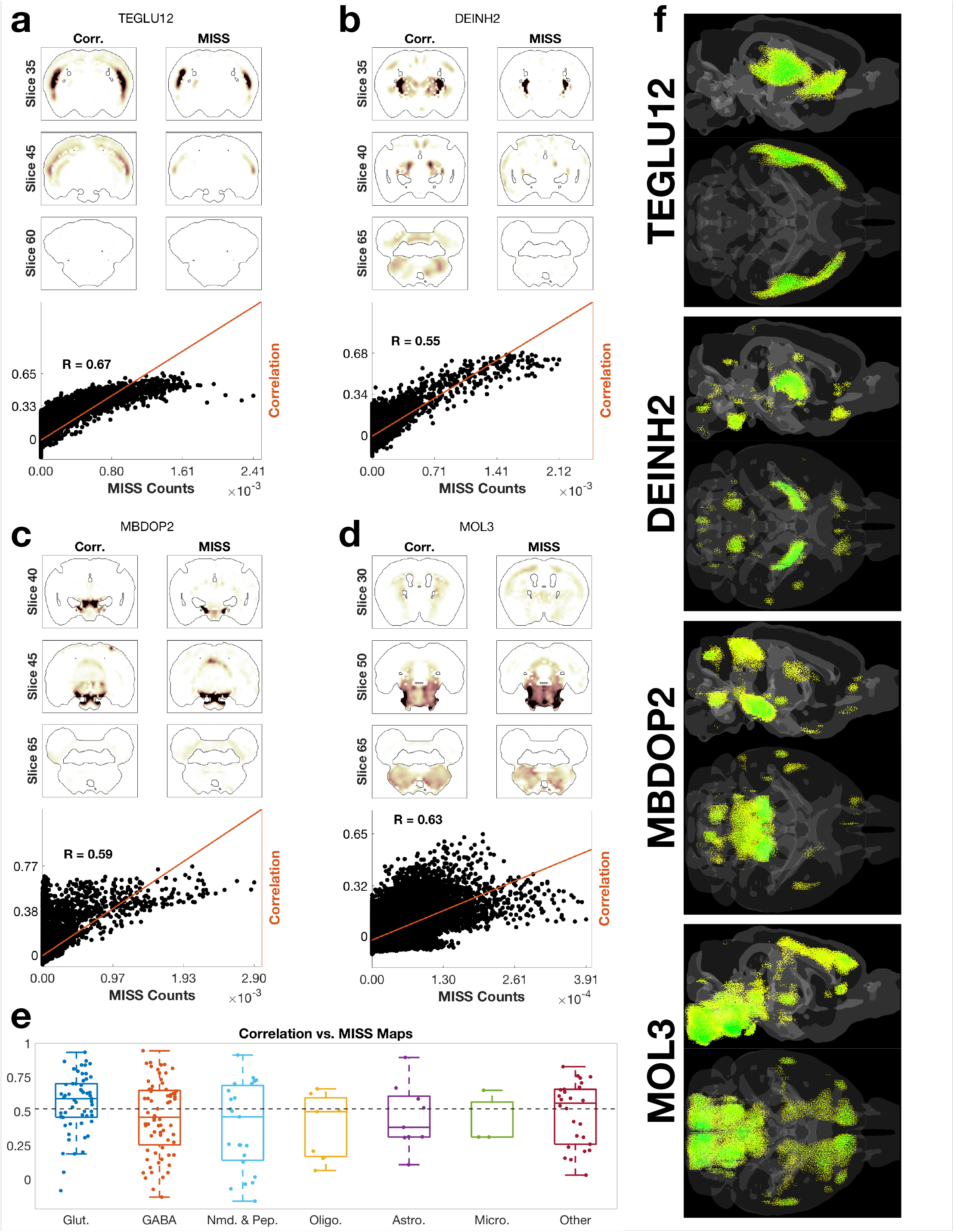
MISS maps of cell classes using data from a more widely sampled scRNAseq dataset with 200 cell classes sampled from a more comprehensive set of brain regions (Zeisel et al., 2018) produce maps that agree with those produced by the original authors. We show comparisons between the two approaches for four distinct cell types: (a) TEGLU12 (telencephalic glutamatergic neuron type 12), (b) DEINH2 (diencephalic inhibitory neuron type 2), (c) MBDOP2 (midbrain dopaminergic neuron type 2), and (d) MOL3 (MOL3-enriched oligodendrocytes). There is strong visual agreement between the approach taken by Zeisel *et al*. and MISS throughout the brain (left and right columns per panel, respectively), as well as strong quantitative agreement at a per-voxel level (scatterplots). (e) Box plots of correlations between the two approaches per cell type, grouped by major cell class. The overall mean and median R-values across all 200 cell classes from Zeisel *et al*. were 0.54 and 0.56, respectively. (f) Axial and sagittal views of the MISS maps of each of the cell types in panels (b)-(e).

## DISCUSSION

### Summary of key results

We provide a method to accurately infer the per-voxel density of a diverse range of neuronal and non-neuronal cell types from gene expression data at a sub-millimeter scale, at both whole neocortical and whole brain levels of coverage. We are able to obtain and evaluate the accuracy of our maps for two key reasons. First, MISS incorporates gene subset selection as a novel and essential preprocessing step, distinguishing it from previous deconvolution approaches for the purpose of mapping cell types (Andersson et al., 2020; Grange et al., 2014). We proposed and thoroughly evaluated a novel subset selection algorithm, MRx3, which outperformed conventional approaches that utilized all available genes. Second, we created novel evaluation metrics for cell type maps and show that our inferred maps give strong quantitative agreement with independent literature-derived regional estimates of GABAergic interneurons (Kim et al., 2017) (**Figure 2**) and faithfully reproduce the laminar architecture of the neocortex (**Figure 3**). We also demonstrate that MISS can be applied to larger scRNAseq datasets with larger numbers of more finely specified cell types (**Figure 5**), generalizing our methodology to other gene-expression datasets.

### Why does MRx3-based subset selection provide the best results?

Our results depend critically on the quality of gene subset selection. This was a novel combination of methodologies, as prior subset selection approaches focused on differential expression or using literature derived marker genes (Beier et al., 2015; Björklund and Dunnett, 2007; Erö et al., 2018; Pazos et al., 1985; Vincent and Kimura, 1992; Zeisel et al., 2018), and prior mapping attempts using deconvolution or matrix inversion did not employ feature selection (Andersson et al., 2020; Grange et al., 2014). Although previous work has suggested that using all available genes provides good mapping results (Andersson et al., 2020; Grange et al., 2014), we find that without filtering of low-information genes, the resulting maps become qualitatively and quantitatively inaccurate in places (**Figures 2 & 3**), with significant diffuse abundance patterns that are biologically implausible (**Figure 3a**; 2^nd^ column). The issue is exacerbated by unsampled cell types in the AIBS data used here (AIBS, 2018; Tasic et al., 2018), since a matrix with missing cell types in **Equation 4** may potentially lead to error in the least-squares solution, particularly in regions anatomically dissimilar to those sampled. Our subset selection step helps ensure that unsampled cell types do not appreciably contaminate the inference of sampled ones, as it specifically selects only the genes most relevant to cells in the scRNAseq datasets. Additionally, many genes are neither specific to the central nervous system, nor do they show appreciable gradients across the brain. The inclusion of such genes can lead to diffuse effects in inferred maps without contributing any useful signal. However, we note that driving feature selection to an extreme is also suboptimal. Small numbers of genes do not give a good performance in R or *τ*_adj_ values (**Figure 3b**), and there is apparent strong cross cell type expression of some genes in the scRNAseq data (**S. Figure 3**; gene names in **S. Table 3**), indicating that the best performance range is achieved at an intermediate subset of genes, whose identification is not trivial.

There are several reasons why MRx3 in particular may yield strong results. While feature selection approaches generally rely upon a measure of differential expression as a criterion for selecting high-information genes, there are other important considerations for determining an optimal gene set for the purposes of matrix inversion-based inference of cell counts. High-information genes are not homogeneously distributed between cell types, meaning that a simple filtering the yields one or a small number of genes per cell type does not perform well (**Figure 3b**). The minimum redundancy criterion contained within the mRMR (Peng et al., 2005) and MRx3 algorithms circumvents this issue by preventing genes from being added if their expression profiles between cell types are too similar to other genes already selected. MRx3 goes a step further than mRMR by also including a minimum residual criterion that prevents genes that, if added, would result in high degrees of error when reconstructing the spatial gene expression matrix from the scRNAseq data and inferred cell type densities. Such genes may be highly noisy in the ISH expression atlas even if they have high information content in the scRNAseq dataset, and will therefore lead to unstable or inaccurate results after matrix inversion (note the sharp increase in residual as those high noise genes are added back into the inversion near the end of the curve in **Figure 3b**). The net effect of removing them, as MRx3 does, is to produce high information gene sets *specifically for the purpose of generating quantitatively validated cell type maps*.

### Further uses and potential applications of the MISS pipeline

Key advantages of MISS are its flexibility and low input cost to generate results. First, MISS is computationally inexpensive and fast, as it performs linear inference on metadata, rather than a time and labor intensive microscopy and image processing pipeline. This allows us to run the entire pipeline from start to finish to generate our cell type maps, including visualization, in hours on a standard laptop. The method achieves strong fits to empirical data despite the fact that choosing the elbow *n*_G_ is dependent only on the input datasets; furthermore, this elbow falls in a range of possible *n*_G_ values that yield quantitatively strong results. We therefore anticipate that as more scRNAseq data becomes available, users will be able to implement the MISS pipeline directly to generate whole-brain maps of yet more cell types; we make all code publicly available on GitHub. Future work includes utilizing the MISS pipeline to infer cell densities in the human brain, using human scRNAseq datasets mapped onto the AIBS human gene expression atlas (Hawrylycz et al., 2012). A computational approach for cell type mapping is particularly appealing in humans, where brain size and tissue accessibility make experimental techniques pioneered in mice prohibitively expensive and time-consuming.

Inferred cell type maps from MISS can also help address the extent to which cellular identities of brain regions governs the formation of synaptic connections (Meyer et al., 2010) and inter-regional neural connectivity (Jones and Rakic, 2010; Szentágothai, 1975). Questions surrounding whether certain behavioral, cognitive, or sensory processing abilities are correlated with certain cell types, their spatial distribution, or their location within connectivity networks (Pinto and Dan, 2015; Roseberry et al., 2016; Senzai and Buzsáki, 2017; Sippy et al., 2015) can also benefit from our maps. Clinically, MISS could further understanding of the selective vulnerability of brain regions such as the entorhinal cortex to early tau inclusions in Alzheimer’s Disease (Braak and Braak, 1991) or the substantia nigra pars compacta to early synuclein inclusions in Parkinson’s Disease (Braak et al., 2003). The spatially varying abundances of cell types considered selectively vulnerable to tau or α-synuclein inclusions can be mapped using MISS, and their correspondence with the spatial pattern of protein pathologies can be tested; for instance, recent experiments suggest cell type selectivity of tau pathology (Fu et al., 2018, 2019; Muratore et al., 2017). Cellular vulnerability in other neurological conditions can also be interrogated using MISS, including psychiatric diseases such as schizophrenia (Skene et al., 2018), and traumatic brain injury, which was hypothesized to preferentially involve certain types of cells in both injury and recovery phases (Arneson et al., 2018; Rama Rao et al., 2018).

### Methodological limitations of the MISS pipeline

The most significant limitation is that the scales of cell densities presented in **S. Table 4** are reliable across voxels, but are not fully so across cell types; it is possible that a per cell type scaling factor could address this issue. In future work, we plan to utilize Nissl and DAPI stains co-registered to the mouse Common Coordinate Framework (CCF) using the connectivity atlas parcellation (Oh et al., 2014) to infer actual counts per voxel. There is also a risk of off-target predictions of cell types due to two confounding factors. First, our method cannot differentiate between gene expression signal in the ISH atlas coming from somatodendritic compartments of cells as opposed to their axon terminals. Second, as not all cell types will be contained in any single scRNAseq dataset, cells not included in a dataset but which have similar gene expression profiles to an included cell could be erroneously mapped. Finally, there is a risk of both overfitting and of choosing a suboptimal value given our selection procedure. However, our selection procedure chooses a value in the middle of a range of values that produce high correlation values between proposed interneuron maps and empirical density data in the neocortex (**Figure 2**) as well as faithful recreations of the laminar patterns of neocortical projection neurons (**Figure 3**). Furthermore, we find no bias towards higher residuals in sampled versus unsampled regions, especially within the neocortex (**S. Figure 2a**). The strength of our results indicates that, despite these limitations, we can reproduce cell densities at per voxel resolution using MISS with superior accuracy.

## Conclusions

We propose a novel computational pipeline for high-accuracy, per-voxel cell type density inference using ISH and scRNAseq data across the entire mouse brain. Our results demonstrate that verifiable mapping of neuronal and glial subpopulations with well-differentiated glutamatergic and GABAergic subpopulations can be obtained using relatively small numbers of cell types and sampled brain regions. Most importantly, we demonstrate that of data-driven gene subset selection prior to cell type mapping, which we accomplish with our novel MRx3 algorithm, is vital for producing more accurate maps. Furthermore, we find that matrix inversion is superior to correlation-based mapping procedures for yielding accurate cell type distributions, but that subset selection was independently important for improving cell type map accuracy regardless of mapping method. We also show that MISS can be applied to other mouse scRNAseq datasets, demonstrating the generalizability of the pipeline. The presented maps and computational pipeline can be used as an inexpensive alternative to single-cell counting for generating density distributions of more cell types than current whole-brain approaches can readily accommodate.

## ACKNOWLEDGEMENTS

The authors would like to acknowledge Dr. Pablo Damasceno for his help with subset selection algorithms and would like to acknowledge Chuying Xia for her prior work in the lab organizing and formatting the raw Allen Institute Mouse Gene Expression Atlas ISH data. The authors would also like to acknowledge Eric Markley for his help conceptualizing the use of an elbow curve for gene set selection.

## AUTHOR CONTRIBUTIONS

Conceptualization, C.M. & J.T., & A.R.; Methodology, C.M., J.T., P.D.M., & A.R.; Writing – Original Draft, C.M. & J.T.; Writing – Review & Editing, C.M., J.T., P.D.M., & A.R.; Figure Generation – C.M. & J.T.; Funding Acquisition, A.R.; Supervision, A.R.

## STAR METHODS

### Input datasets

Input data for cell type or class density inferences were derived from publicly available murine datasets, including the Allen Gene Expression Atlas (AGEA) from the Allen Institute for Brain Science (AIBS). We include a summary of the methodologies of these datasets below in the “Single-cell RNA sequencing” and “In-Situ Hybridization (ISH)” subsections, but as we did not perform these experiments, methodological details should be found in Tasic *et al*., 2018, Zeisel *et al*., 2018, Lein *et al*., 2007, and on the AIBS website.

#### Single-cell RNA sequencing

We utilized all currently available single-cell RNA sequencing (scRNAseq) data from the AIBS website (http://celltypes.brain-map.org/rnaseq), consisting of 27477 individual cells isolated from three distinct brain regions (primary visual cortex (VISp), 15413 cells; anterior lateral motor cortex (ALM), 10068 cells; lateral geniculate complex (LGd); 1996 cells) profiled across 45768 unique genes. The cells in the database were then classified into unique clusters, which were taken to be distinct cell types, based on differential expression scores of a reduced set of differentially expressed genes. The authors implemented a multistep clustering pipeline that first utilizes two complementary dimensionality-reduction algorithms to reduce the sample space of the data, iterative principal component analysis (PCA) and iterative weighted gene co-expression network analysis (WGCNA), then clusters the resulting “modules” hierarchically using the Jaccard-Louvian method (Shekhar et al., 2016) or Ward’s method, depending on the number of cells to be clustered. The resulting clusters were identified and validated using three sources of independent information: the layer-enriching dissection from which the cells were extracted, the presence of differential expression markers corresponding to known or newly discovered cell types, and the localization of these markers as determined by *in situ* hybridization. Based on these labels, the clusters were further grouped into a three-level hierarchy, representing broader classes of cell types that were of particular interest in the present study. The high robustness of this labeling and classification of cell types within this particular scRNAseq dataset allowed us to confidently determine neocortical spatial patterns, since the cell type labels themselves were already validated. For further methodological details, see Tasic *et al*. (Tasic et al., 2018) and the RNAseq documentation on the AIBS website (http://celltypes.brain-map.org/rnaseq).

We also mapped CNS cell data available through the Mouse Brain Atlas (MBA; http://mousebrain.org), as the sampling of types across the entire cortex was more comprehensive (Zeisel et al., 2018). Low-quality samples were removed and clustering into types was performed through a novel multilevel approach employing PCA, k-nearest neighbors (KNN), and t-distributed stochastic neighbor embedding (t-SNE) on a reduced, high-information gene set determined through a support vector machine (SVM) classifier. The final dataset contains 265 unique cell types, of which we have mapped 200 (excluding types specific to the spinal cord, which is not present in the ISH atlas); in all, we incorporated data from 144147 individual cells sampled across 12 major region groups. For further methodological details, see Zeisel *et al*. (Zeisel et al., 2018) and the RNAseq documentation on the MBA website (http://mousebrain.org).

#### Normalization of Input scRNAseq Datasets (Tasic *et al*.)

The AIBS scRNAseq data exists as tables of raw exon and intron counts (sequence hits) separated by sample ID and by gene for three distinct brain regions. For each of these regions, we combined the raw exon and intron counts into a single metric according to AIBS guidelines:

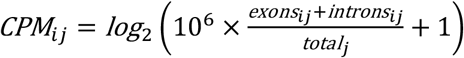

where CPM_*ij*_ is the sample-normalized expression (in counts per million) of gene *i* in sample *j* and total_*j*_ is the total number of reads for all genes in sample *j*. We then utilized the existing cluster identification for each sample (excluding those labeled “low quality”, “doublet”, or “outlier”) to derive each gene’s mean expression for each cell type previously determined in the AIBS datasets (AIBS, 2018; Tasic et al., 2018). For samples extracted from neocortical regions (Tasic et al., 2018), we used the correlation coefficients between their individual expression and their cluster’s center-of-mass to infer a more accurate mean expression for the cluster; for samples from the lateral geniculate complex (AIBS, 2018), no such information was available and an unweighted average was used instead. The resulting clusters, however, represent cell types that were too specific for our purposes, as we were interested in the spatial localization of the broader subclasses and classes of cell types (e.g. all Layer-5 pyramidal tract neurons as a single group). Some rarer types also contained very few individual cells. We therefore determined the expression of these broader cell type designations by averaging the expression of the clusters assigned to them across sampled regions, weighted by the number of samples per cluster; it has been previously shown that when cell types are combined in this way, under matrix-inversion approaches the resulting inferred counts represent a combined signal of the constituent cell types (Grange et al., 2014). The resulting dataset, herein referred to as the “Tasic *et al*. dataset,” contains genome-wide expression data for 21 neuronal and 4 non-neuronal cell types (**S. Figure 1a, S. Table 1**). More specifically, the following groups of cell types are represented: 1) neocortically derived, layer-specific glutamatergic neurons; 2) neocortically derived GABAergic interneurons (*Pvalb*+, *Sst*+, and *Vip*+); 3) four non-neuronal types (microglia, astrocytes, oligodendrocytes, endothelial), all derived from the neocortical and thalamic regions; and 4) various other glutamatergic and GABAergic neurons which were labeled with marker names we could not trace back to well-characterized cell types. Several of the non-neuronal cells represent a pooled signal between related, associated types because of the relative scarcity of these types in the dataset: “microglia” represents a combined signal across specific types of microglia and macrophages, “oligodendrocytes” represents a combined signal across specific types of oligodendrocytes and OPCs, and “endothelial” represents a combined signal across true endothelial cells, pericytes, SMCs, and other vasculature-related types.

#### Normalization of Input scRNAseq Datasets (Zeisel *et al*.)

We followed a similar procedure as above to obtain the input scRNAseq dataset derived from the MBA database (http://mousebrain.org). The expression data were already log2-normalized by Zeisel *et al*.; therefore, the only pre-processing necessary prior to MISS was excluding non-CNS cells, leaving us with a dataset (herein referred to as the “Zeisel *et al*. dataset”) comprised of 200 types. Unlike with the Tasic *et al*. dataset, we chose to map each without combining them into larger clusters.

#### *In situ* hybridization (ISH)

Our spatially realized expression data were derived from the Allen Gene Expression Atlas (AGEA), a publicly available ISH dataset published by the AIBS (https://mouse.brain-map.org/search/index). We specifically utilized the coronal dataset, which contains better whole-brain spatial coverage than the sagittal series, at the cost of less genomic coverage (4344 ISH probes covering 4083 unique transcripts). The brains of 56-day-old C57BL/6J mice were extracted and sectioned in 25-μm slices, which were separated out into 8 series of 56 slices each uniformly spaced 200-μm apart. Each riboprobe corresponded to one series and was designed to have minimal overlap with other transcripts to prevent signal contamination by off-target binding. Post incubation with the probe, slices were imaged in high resolution (1.07-μm pixels) and coregistered onto a common 3-dimensional space. The expression data were downsampled by averaging the grayscale intensity of expressing pixels within a 200-μm by 200-μm area and dividing by the total number of pixels in that area (“expression energy”), yielding a measure of gene expression over a rectangular volume of 67 by 41 by 58 voxels. For further details, see Lein *et al*. (Lein et al., 2007) and the ISH documentation from the AIBS (http://help.brain-map.org/display/mousebrain/Documentation). We normalized each gene’s expression by its sum across all voxels, so that they were on comparable scales for cell count inference.

### MISS Pipeline

#### Cell Density Inference

Our overarching goal in this study was to obtain accurate densities for individual cell types across the whole mouse brain. Grange *et al*. first introduced a mathematical framework for inferring voxel-wise cell type densities, which posits that ISH voxel energy for a given gene is proportional to each cell type’s expression value for that gene multiplied by its density in that voxel, summed across all cell types within that voxel (Grange et al., 2014). Recent similar prior work used scRNAseq profiles to infer cell densities within spatial RNAseq sampled areas (Andersson et al., 2020).

We can generalize the relationship between expression energy per voxel and cell type-specific gene expression across all voxels and genes in matrix form as follows:

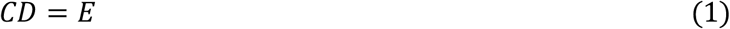

where *E* is the row-normalized genes-by-voxels (*N*_*G*_ × *N*_*V*_) expression matrix extracted from the ISH data, *C* is the row-normalized genes-by-cell types (*N*_*G*_ × *N*_*T*_) expression matrix extracted from the scRNAseq data, and *D* is a cell types-by-voxels (*N*_*T*_ × *N*_*V*_) matrix of cell densities. To solve **Equation 1** the matrix *C* must be inverted in some manner; let us denote for convenience *C*^#^ a suitably defined pseudo-inverse of *C*. Hence we estimate:

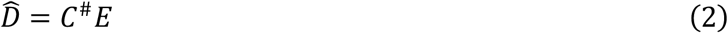

In practice, a classical Moore-Penrose pseudo-inverse of *C* is unsatisfactory since it does not enforce the biological fact that cell densities cannot be negative. Hence in this paper 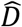 was estimated using the `lsqnonneg` function in MATLAB, a least-squares solver with a built-in non-negativity constraint:

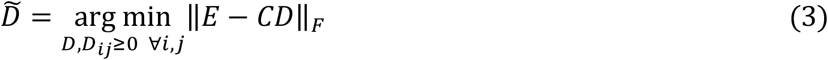

Prior approaches relying upon matrix inversion have utilized all available genes, which could introduce noise into the predicted cell type densities. Here we introduce a novel subset selection procedure, Minimum Redundancy – Maximum Relevance – Minimum Residual (MRx3), which ranks genes for inclusion by information content and then excludes all genes that fall below a certain threshold prior to inversion (**Figure 1, Steps 2 & 3**). The MISS algorithm first creates reduced matrices *C*_*red*_ and *E*_*red*_, which contain only MRx3-selected genes, and then solves the non-negative least-squares problem posed above, substituting *C*_*red*_ and *E*_*red*_ for *C* and *E*, respectively, in **Equation 3**. The procedure for MRx3 subset selection is detailed in the following section.

We then directly compared the resulting inferred densities, averaged across anatomical regions, to those determined in previously published work (Kim, et al., 2017), and derived novel metrics to further assess the accuracy of our mapping (See **Method Validation** below). Of note, a potential source of error in the inversion is that not all cell types in the Tasic *et al*. dataset contain the same total number of transcripts due to differences in cell size and developmental origin. However, the variance between the number of unique gene expressed per cell type that are present within the AIBS gene expression atlas is low, mitigating this source of potential error (**S. Figure 4**).

#### Gene subset selection

A fundamental limitation to the matrix inversion scheme (**Equation 1**) is that it implicitly assumes that the columns of *C* span all of the relevant cell types in all voxels across the brain. This assumption has underlain the approach of prior inversion or deconvolution based cell type mapping pipelines. In our case, where we derive *C* from scRNAseq data obtained from only three distinct anatomical regions, this assumption is clearly violated. Furthermore, many genes carry only limited information in differentiating cell types represented in our dataset, and therefore only add noise to the system. Both problems can be ameliorated by performing the matrix inversion only over the set of maximally informative genes. This is accomplished by removing rows of *C* and *E* corresponding to low-information genes and forming reduced-dimension versions: *C*_*red*_ and *E*_*red*_. Determining such a subset is a nontrivial optimization problem, both in terms of reordering genes by information content and finding the optimal subset size. To solve the first problem, we introduce the MRx3 algorithm, which we describe in detail below (see also **Figure 1, Step 2**). We then calculate the Frobenius norm of the residual between the true spatial gene expression and the cell-density-imputed spatial gene expression, ‖*E*_*red*_ − *C*_*red*_ · *D*‖_*F*_, for a range of subset sizes and choose an optimal subset by finding the elbow of the resulting L-curve (**Figure 1, Step 3**). This elbow, which we define as the point closest to the origin, is the subset that optimizes the tradeoff between residual and number of genes used. We applied this elbow selection procedure to the Tasic *et al*. and Zeisel *et al*. scRNAseq datasets independently, since gene information content and subset size are highly dependent on the set of cell types contained within each.

#### MRx3 Algorithm

In the current work we propose the following Minimum Redundancy – Maximum Relevance – Minimum Residual (MRx3) gene subset selection algorithm, which is based upon the popular mRMR algorithm. The mRMR algorithm attempts to identify the maximally informative set of features with minimal overlap between features, and has been successfully applied to microarray data (Peng et al., 2005). In brief, it utilizes a simple greedy search approach to iteratively add genes, one at a time, that maximize an objective function defined by each gene’s degree of differential expression across all cell types divided by its similarity to genes that have already been incorporated (this corresponds to the original mRMR algorithm’s “quotient” variant, which was determined to give superior performance (Peng et al., 2005)).

We assume for simplicity that the variance per entry of *C*_*col*_ is uniform. Then we define a suitable F-statistic on our data, for any candidate gene *i* ∈ *G* as:

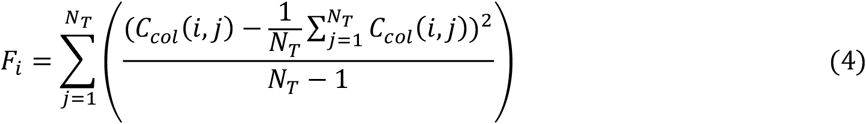

This quantity corresponds to the maximum relevance criterion previously proposed (Peng et al., 2005). The minimum redundancy criterion is given by the mean of the absolute value of the Pearson correlation between the candidate gene *i* and the already-selected genes within *S*:

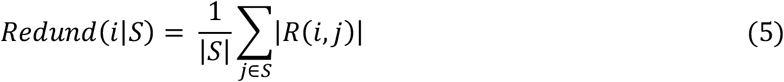

Using these two metrics we implemented the following version of the mRMR algorithm that we applied to the rows of *C*_*col*_.

<u>Algorithm mRMR:</u>

1. Initialize *S*_0_ = ∅
2. For *k* ∈ [1, *n*_*G*_], iterate:

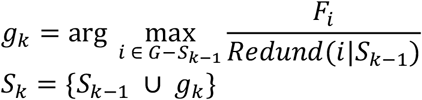

Essentially, this procedure finds, from the currently unselected pool *G* − *S*, a single candidate gene per iteration that maximizes the F-statistic while minimizing its correlation with those genes that are already in the selected set *S*. Using the selected genes we finally create row-decimated matrices

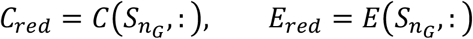

Although the mRMR approach outlined above produces gene sets that appropriately capture cell type-specific features, it does not consider the impact of these genes on the overall prediction error between *E*_*red*_ and *C*_*red*_*D*. The original mRMR algorithm was developed in the context of classification, where the two criteria of relevance and redundancy are appropriate. Here, we have a linear regression problem given by **Equation 1**, hence we have in addition a third criterion: *that the reduced matrix produce low residuals – this is the* MRx3 *approach*. This term should penalize genes for adding exorbitant prediction error, as we anticipated that these genes, while maximally relevant and minimally redundant with respect to the scRNAseq dataset, would negatively affect the accuracy of the resulting inferred *D*.

We therefore calculate the projection error each gene contributes to the overall projection error in the inference described in **Equation 1** using rank-1 update rule. For every candidate gene, *g*_*i*_, the complementary gene set of size *N*_*g*_ − 1 (denoted by the subscript ∼*i*) can be constructed, which contains all other genes. The spatial gene expression residual for this complementary set can be analytically evaluated using the pseudoinverse: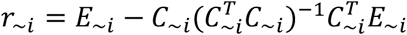. To then find the individual contribution of gene *g*_*i*_, we augment the current gene set with a new candidate gene *g*_*i*_, such that 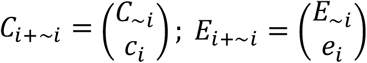 with *c*_*i*_ = *C*(*g*_*i*_, :) and *e*_*i*_ = *E*(*g*_*i*_, :), the following recursion equation can be demonstrated:

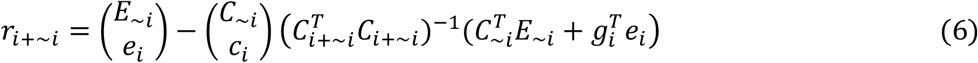

Now, *C*_*i*+∼*i*_ has one row more than *C*_∼*i*_, hence to evaluate the matrix inverse above, we use the Sherman-Morrison rank-1 update formula (Sherman and Morrison, 1950), and assert that 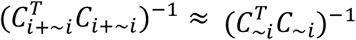, which is good for ‖*C*_∼*i*_*c*_*i*_‖ ≪ 1. The approximation is further improved for a sufficiently large selected set. In this event, the squared prediction error *ϵ*_∼*i*_= ||*r*_∼*i*_ | | _*F*_ ^2^ accommodates the following update rule for gene set *S*_*i*+∼*i*_ = *S*_∼*i*_ ∪ *g*_*i*_:

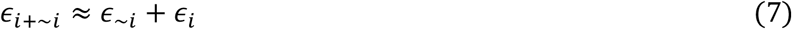

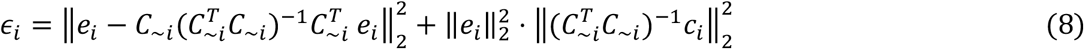

Note, the second term in **Equation 8** is an upper bound arising from the Cauchy-Schwartz inequality. This update rule is cheap to evaluate since it does not require computing the matrix inverse from scratch, and only involves matrix multiplication. Once we have calculated the *ϵ*_*i*_ for each candidate gene, we remove all genes with projection error at or above the 90^th^ percentile from the active set, only adding them back after all low-error genes have been reordered by mRMR. This choice accounts for the the rapid increase in residual per gene at the tail end of the curve in **Figure 3b**.

We therefore can denote our mRMR + Minimum Residual, or MRx3 algorithm as follows:

<u>Algorithm (MRx3):</u>

a. Calculate projection error per gene, as denoted in **Equation 8**, and exclude the genes at or above the 90^th^ percentile of projection error.
b. Run <u>Algorithm mRMR</u> until all genes not in the excluded 10% based on our projection error metric are ranked and ordered.
c. If *n*_*G*_ > 0.9 * *N*_*G*_, add the remaining 10% of excluded genes back in one by one.

Here the percentile set is always the 90^th^ percentile, or the top 10% of genes. While this could be set as a tuneable parameter, given the behavior of the residual in the elbow curves (**Figure 1, Step 3**; **Figure 3b**), we believe this is unnecessary in this case.

## Method Validation & Comparison

### Direct comparisons to literature

We utilized existing knowledge about cell counts in the mouse brain where they have been established. Three recent datasets, which we chose for their broad spatial coverage, include quantifications of three distinct classes of GABAergic neurons (Kim et al., 2017). From these literature-derived cell counts, we create regional densities of each interneuron class by dividing the counts by the regional volumes. We then averaged across all voxels in each region in the AGEA for each of these three GABAergic interneuron classes in our cell type maps. We limited these comparisons to neocortical regions, as the classes of GABAergic interneurons whose scRNAseq profiles were used for mapping in this paper were sampled from two neocortical regions (Tasic et al., 2018), making the accuracy and interpretability of maps beyond the neocortex uncertain.

We also directly compared our MISS estimates on an independent, and larger scRNAseq dataset from a wider array of sampled brain regions (Zeisel et al., 2018) with maps using the correlation-based mapping procedure and gene sets from that same publication. Particularly, we assessed how correlated, on a per voxel level, the proposed R-values (from correlation-based mapping) and densities (from MISS) of each cell class were. Further information on the correlation-based mapping procedure can be found in the Correlation-Based Mapping subsection.

### Layer ordering

Seven of the twenty-five cell types represented within our sample are excitatory neurons which have been specifically extracted from individual neocortical layers (Tasic et al., 2018) (**S. Table 1**). Although we could not find corresponding literature estimates at this level of cell type specificity, we expected that we could accurately recreate the appropriate layer ordering of the neocortex from these cell types, because they have mostly non-overlapping distributions at the spatial resolution of the ISH dataset. Moreover, such an ordering would demonstrate that our inference is accurate not only at the level of anatomical regions, but sub-millimeter voxels as well. We implement the following procedure to infer the neocortical layer ordering per coronal slice:

1. Initialize *y, z* = 0.
2. Map the per-voxel cell counts of each layer-specific glutamatergic neuron across all slices of the brain volume and apply modest smoothing to the resulting images.
3. Determine the center line using MATLAB’s `bwskel` of each layer-specific glutamatergic neuron’s density “band” within the neocortical voxels of a given slice.
4. Denote the number of cell types that have no neocortical signal by *y*′ and the number that has more subcortical, midbrain, and hindbrain signal than neocortical signal by *z*′. Let *y* = *y* + *y*′ and *z* = *z* + *z*′.
5. Calculate the mean distance of each center line to the cortical surface and rank the cell types from closest to farthest.
6. Repeat Steps 2 through 5 for each coronal slice containing neocortical voxels and tabulate the results in a cell types-by-slices ordering matrix *O*.

For each column (slice) of *O*, we produced a corresponding column in the ground-truth ordering matrix *O*^*^, which uses the glutamatergic cell type labels to form the expected layer order among cell types that were present in that slice. Since these orderings contain many identical entries, we utilized a nonparametric rank correlation that accommodates ties, Kendall’s *τ*, between the columns of *O* and the columns of *O*^*^ vertically concatenated. Since we expected these layer-specific neurons to be expressed throughout the neocortex, we incorporated a sensitivity and specificity penalties through *y* and *z*, respectively, and calculated the following adjusted correlation:

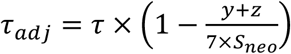

where *s*_*neo*_ is the number of coronal slices containing neocortex. We calculate this value for our optimal MISS estimates, matrix inversion with no subset selection, and correlation-based mapping.

### Correlation-Based Mapping

As a point of comparison with inversion and deconvolution based mapping approaches, we also adopted a correlation-based mapping procedure similar to the one enumerated in Zeisel *et al*. (Zeisel et al., 2018). This procedure calculates the correlation between the Z-scores of each gene across cell types or classes in scRNAseq space, *Z(C)* versus the Z-score of each gene across voxels in ISH space, *Z(E)*. The algorithm can be described as follows:

1. Calculate the Z score per gene across cell types of classes in scRNAseq space, *Z(C)*
2. Calculate the Z score per gene across voxels in ISH space, *Z(E)*
3. Correlate *Z(C)* with *Z(E)* to get an estimate per voxel of the correlation between each cell type and each voxel in terms of per gene Z scores using Pearson Correlation
4. Set all R-values below 0 to 0

Further methodological details can be found in the original publication (Zeisel et al., 2018).

### 3D Visualization of Cell Type Distributions

Our whole brain illustrations of MISS-inferred cell type distributions are rendered using an in-house brain visualization package. We first upsample and co-register our inferred cell type maps, which inherits the 200μm resolution from the AGEA (Lein et al., 2007), to the 100μm AIBS mouse CCF atlas (AIBS, 2013; Oh et al., 2014). We then use a point cloud overlaid on a surface rendering of the mouse brain volume to illustrate the distributions of each cell type across the whole brain. Importantly, each point does not represent the exact location of an individual cell. Instead, the number of points is controlled per voxel by the density of each type of cell in each voxel. The location of each point is given by voxel location, with slight random jitter added to each point to prevent complete overlap, which would obscure the visualization. We emphasize that these images are not, and should not be interpreted as, providing the exact location of any individual cells.

### Robustness Analyses

We performed two robustness checks, shown in **S. Figure 2**, to examine whether our MISS-inferred cell type maps are robust to noisy data, perturbation, and the sparse sampling of regions in the scRNAseq dataset. First, we calculated the per-voxel residual between the scRNAseq data and the AGEA (Lein et al., 2007) across both scRNAseq-sampled and unsampled regions and compared the distribution of residuals across both of these sets of voxels (**S. Figure 2a**). We also averaged per-voxel residuals into brain regions and mapped these residuals to verify sampled regions, at the overall regional level, did not show different residuals than other regions (**S. Figure 2b**). Finally, we also demonstrate high correlation between the cell types mapped using the gene set given by the chosen elbow and those gene sets with up to 100 fewer or more genes included, with inclusion given by MRx3 ranking (**S. Figure 2c**). We further highlight that the plotted gene set is within a range of higher quality gene sets (for mapping purposes) in **Figure 3b**, as we demonstrate that the gene set given by the elbow index is around the middle of a range of genes that produce strong R-values compared with empirical neocortical GABAergic interneuron densities and recreates expected laminar order of neocortical glutamatergic neurons.

### Implementation

The MISS pipeline is written in version 2020a of MATLAB and is publicly hosted on GitHub (https://github.com/Raj-Lab-UCSF/MISS-Pipeline/). All data processing, analysis, and visualization were run on a MacBook Pro with 16 GB of RAM and a 3.1 GHz Quad-Core Intel Core i7, although we anticipate that any machine capable of running the current version of MATLAB with at least 8 GB of RAM can successfully run the MISS pipeline.

---

**S. Figure 1.**
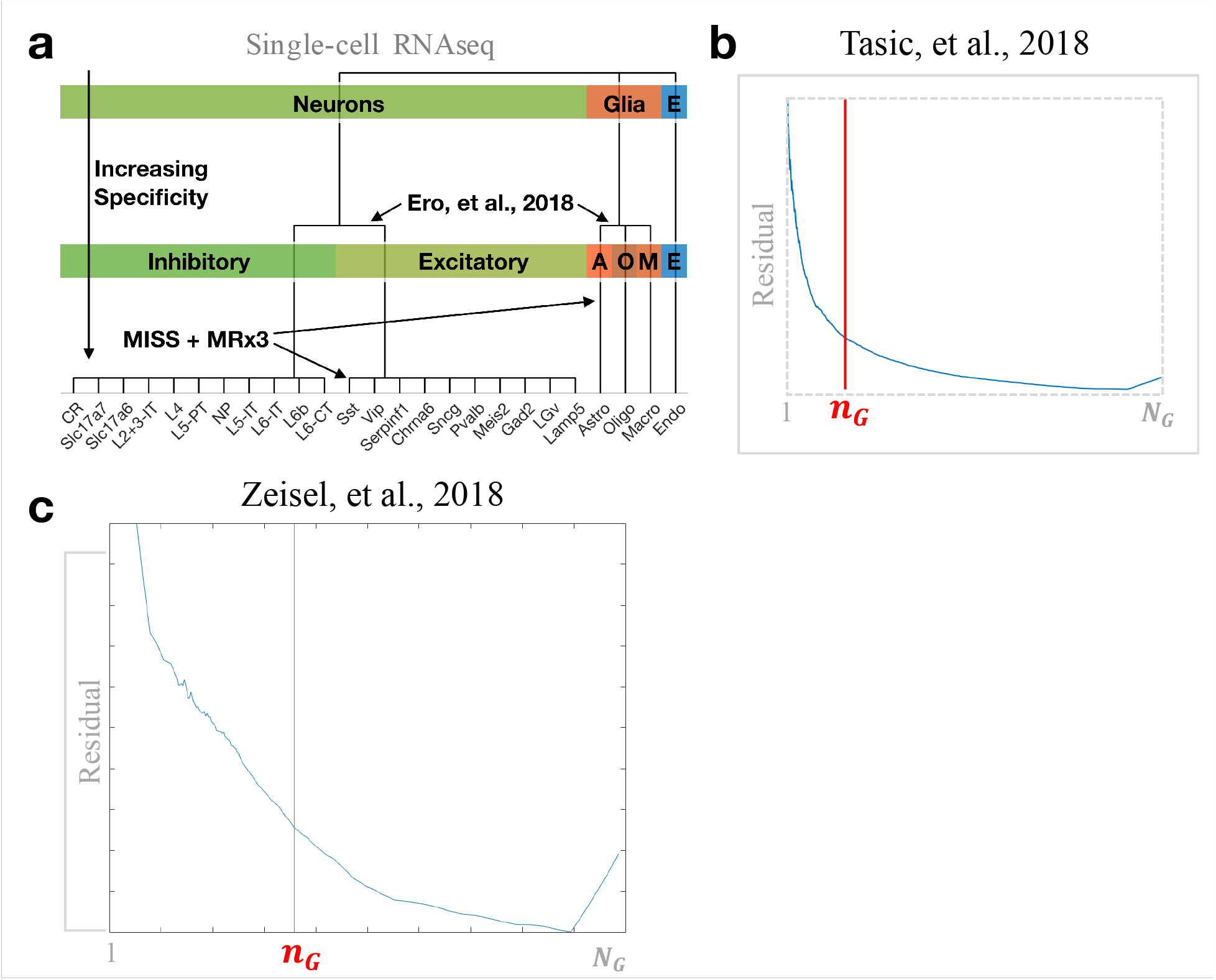
Cell class level hierarchy & elbow gene index selection before cell type mapping. (a) Here we demonstrate that the scRNAseq data from Tasic *et al*. (AIBS, 2018; Tasic et al., 2018) is hierarchically clustered, and we show the level, including cell type/major cell class names that we chose to map; the rationale for mapping at the selected level of hierarchy is because neuronal cell types such as *Pvalb*+, *Sst*+, and *Vip*+ GABAergic interneurons, as well as glutamatergic laminar projections neurons from the neocortex, such as L6-CT or L4 neurons, are clear, well-accepted cell class labels whose resultant maps could be compared with empirical data and a priori expectations, aiding validation of the method. We show the elbow selection curve, with elbow defined as the point closest to the origin on the curve of L1-normalized MRx3 gene rank versus ||*E* − *CD*||_*F*_ for mapping both (b) the scRNAseq profiles from Tasic *et al*. at *n*_G_ = 606, and (c) the scRNAseq profiles from Zeisel *et al*. (Zeisel et al., 2018) at *n*_G_ = 1360.

**S. Figure 2.**
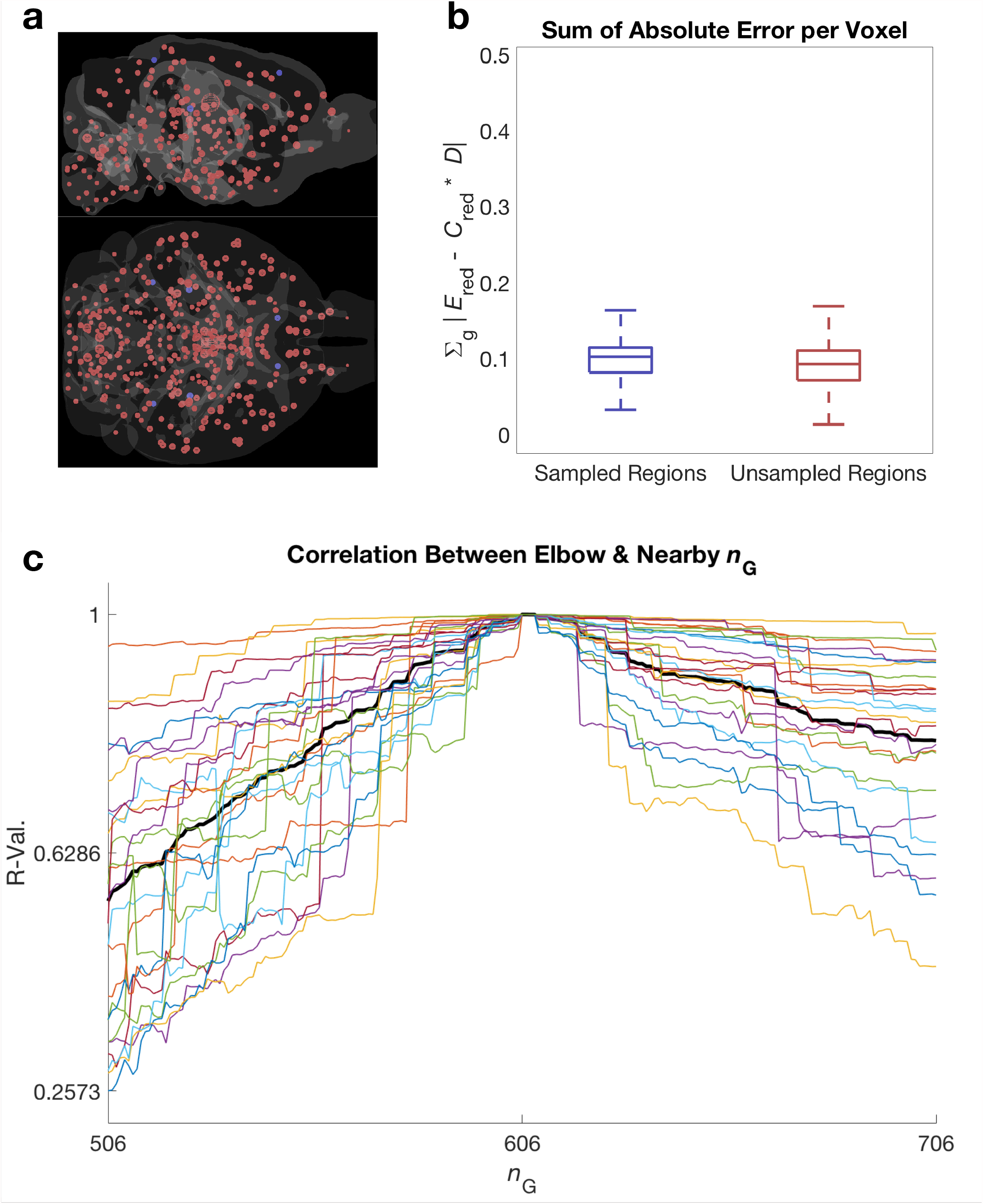
Matrix inversion shows no residual bias towards sampled regions and maps produced with gene indices near the selected elbow show strong per-voxel similarity across cell types to maps produced using the elbow gene index. We show both visually in a 3D mouse brain volume in sagittal and axial views (a), as well as in a box and whisker plot (b), that regions from which the scRNAseq data in Tasic *et al*. were sampled from do not show a lower residual after inversion than unsampled regions. In these plots sampled regions are colored blue and unsampled regions are colored red. (c) Within 25 genes ranked on either side of the chosen elbow gene, no mapped cell type using the scRNAseq profiles from Tasic *et al*. is correlated less than 0.7 at the per voxel level, with the average correlation being close to 0.9. By 50 genes ranked on either side of the chosen elbow gene, the average correlation drops to just above 0.85, but no cell type map is correlated with elbow at less than 0.5. We only see a major drop-off as we go toward 100 gene ranks away from the chosen elbow gene on either side.

**S. Figure 3.**
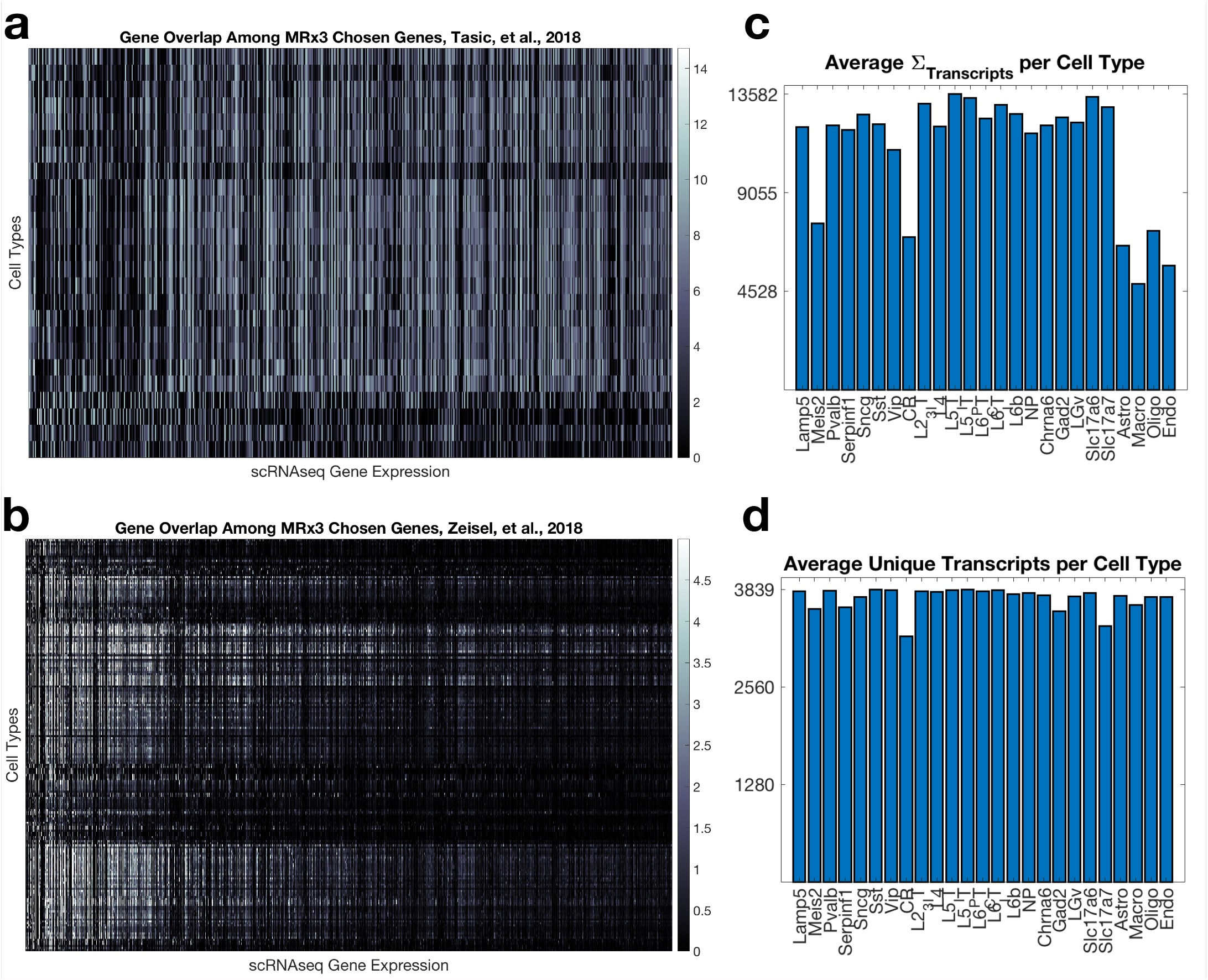
Most genes in the MRx3 subset are specific to one or a small handful of cell types, but some show nontrivial overlap across many types, while the labeled cell types/classes from the scRNAseq data in Tasic *et al*. show differences in total transcript count, but none in number of unique transcripts. Among the genes chosen by MRx3, both those from (a) Tasic *et al*. and (b) Zeisel *et al*., there is generally little overlap between cell types in terms of expression. However, some chosen genes show wide expression across cell types, demonstrating that selection of these MRx3 sets, and cell type mapping gene sets in general, is not necessarily trivial. (c) Cell classes or types from the scRNAseq data from Tasic *et al*. show differences in total transcript count, but not (d) in total number of unique transcripts expressed per type.

**S. Table 1**. We provide a list of major classes of neuronal and glial cell types from the Tasic *et al*. dataset (AIBS, 2018; Tasic et al., 2018) with a metric of whether the type of cell is definitively a canonically well-characterized cell type. We note that *Pvalb*+, *Sst*+, and *Vip*+ GABAergic interneurons, the various laminar neocortical glutamatergic neurons, and the glia are well-characterized types by their major class, but other neuronal subtypes are not, necessarily.

**S. Table 2**. AGEA region names, reordered by canonical major region groupings, including amygdala, following traditional anatomical hierarchy and nomenclature (Paxinos and Franklin, 2012). All regions are ordered in this manner across all files and code for standardization.

**S. Table 3**. Gene names for the genes chosen by MRx3 from the unity set between scRNAseq datasets (AIBS, 2018; Tasic et al., 2018; Zeisel et al., 2018) and the spatial ISH expression atlas (Lein et al., 2007). The file is an .xlsx with the first sheet containing the MRx3 subset using the Tasic *et al*. dataset and the second sheet containing the MRx3 subset using the Zeisel *et al*. dataset.

**S. Table 4**. MISS-inferred regional cell type densities of the twenty-five neuronal and glial major clusters mapped using the Tasic *et al*. scRNAseq dataset (AIBS, 2018; Tasic et al., 2018). This file is a .csv with columns representing cell types and rows representing regions.

**S. Table 5**. MISS-inferred regional cell type densities of the 200 neuronal and glial major clusters mapped using the Zeisel *et al*. scRNAseq dataset (Zeisel et al., 2018). This file is a .csv with columns representing cell types and rows representing regions.

