## Supplementary material for "Matrix Inversion and Subset Selection (MISS): A novel pipeline for mapping of diverse cell types across the murine brain": Table S1

| Cell Type Name[1,2] | Major Cell Class | Derived From: | Description |
|---|---|---|---|
| L2/3 IT | Glutamatergic | VISp, ALM | L2/3 intratelencephalic projection neuron |
| L4 | Glutamatergic | VISp, ALM | L4 projection neuron |
| L5 IT | Glutamatergic | VISp, ALM | L5 intratelencephalic projection neuron |
| L5 PT | Glutamatergic | VISp, ALM | L5 pyramidal-tract projection neuron |
| L6 CT | Glutamatergic | VISp, ALM | L6 corticothalamic projection neuron |
| L6 IT | Glutamatergic | VISp, ALM | L6 intratelencephalic projection neuron |
| L6b | Glutamatergic | VISp, ALM | L6b projection neuron |
| NP | Glutamatergic | VISp, ALM | Near-projecting neuron (?) |
| CR | Glutamatergic | VISp, ALM | Cajal-Retzius neuron (?) |
| Slc17a6 | Glutamatergic | LGd | (?) |
| Slc17a7 | Glutamatergic | LGd | (?) |
| Pvalb | GABAergic | VISp, ALM | Pv+ interneuron |
| Sst | GABAergic | VISp, ALM | Sst+ interneuon |
| Vip | GABAergic | VISp, ALM | Vip+ interneuron |
| Serpinf1 | GABAergic | VISp, ALM | Serpinf1+ inhibitory neuron (?) |
| Sncg | GABAergic | VISp, ALM | Sncg+ inhibitory neuron (?) |
| Chrna6 | GABAergic | VISp, ALM | Chrna6+ inhibitory neuron (?) |
| Meis2 | GABAergic | VISp, ALM | Meis2+ inhibitory neuron (?) |
| Lamp5 | GABAergic | VISp, ALM | Lamp5+ inhibitory neuron (?) |
| LGv | GABAergic | LGd | (?) |
| Gad2 | GABAergic | LGd | (?) |
| Astro | Glial | VISp, ALM, LGd | Astrocyte |
| Macro/Micro | Glial | VISp, ALM, LGd | Microglia + macrophage |
| Oligo | Glial | VISp, ALM, LGd | Oligodendrocyte + OPC |
| Endo | Endothelial | VISp, ALM, LGd | Endothelia + other vessel-associated cells |
